# Comparative Evaluation of Conventional Inorganic Fertilization and *Sesbania rostrata* Green Manuring on Soil Properties and the Growth and Development of *Oryza sativa L. ‘Pant Basmati 1’*

**DOI:** 10.64898/2025.12.24.696455

**Authors:** Hem C. Joshi, Babita Patni, S.K. Guru, Manoj Kumar Bhatt, Meenal Singh

**Author notes:** **Corresponding Author:** Hem Ch. Joshi Manoj Kumar Bhatt.

## Abstract

To assess the impacts of conventional and organic nutrient management techniques on growth, productivity, sustainability, and soil health in the Basmati rice (*Oryza sativa* L.) cultivar *Pant Basmati 1*, a two-year field experiment was carried out during the Kharif seasons of 2023 and 2024. Crop duration remained stable across management methods, as illustrated by phenological variables such as panicle initiation (58-62 days), blooming (84–88 days), and maturity (106-111 days) that were not significantly affected by fertilizer source. Conventional fertilizer supported superior growth at later stages, while organic management produced greater plant height at early growth phases. While the leaf area index and dry matter accumulation were consistently higher under conventional management because of the quick availability of nutrients, tiller density remained similar across treatments (363-411 m⁻²). Despite statistically significant variations, chlorophyll a and b quantities peaked under organic inputs at 60 DAT. Higher biological and grain yields were obtained with conventional fertilization; however, yield variations between systems were generally insignificant over time. As soil fertility stabilized, organic management notably increased the harvest index in the second year, indicating improved assimilate partitioning. The findings show that organic nutrient management can provide benefits in terms of resource efficiency and environmental sustainability while maintaining yield parity with conventional systems during the early conversion phase. Organic management techniques improve soil quality. Compared with to conventional methods, organic management practices were found to have higher soil macro and micronutrient status, water-holding capacity, aggregate stability, and soil enzyme activity (DHA and acid & alkaline phosphatase). These results show that organic Basmati rice cultivation is a feasible and sustainable alternative to traditional nutrient management, even though it may take longer to reach maximum yield.

## Introduction

Soil health and quality determine environmental quality (Pierzynski et al., 1994), the sustainability of agriculture (Acton and Gregorich, 1995), and, consequently, human, animal, and plant health (Habern, 1992). The concept of “soil quality” has been used in agriculture to describe a soil’s ability to sustain crop growth without degradation. It is an essential part of sustainable agriculture and comprises the physical, chemical, and biological properties that enable soil to perform various functions. Productivity, diversity, and soil biological activity are crucial soil functions (Karlen et al., 1997). Karlen et al. (1997), soil biological activity, productivity, and diversity are essential soil functions. The quantity and quality of organic carbon, which determine biological traits substantially more than chemical and physical factors, are the primary factors influencing soil quality. Soil quality indicators are strongly influenced by various agricultural and plant systems and management methods (Tolanur & Badanur, 2003; Ali et al., 2025a; Shagun et al., 2024).

Despite stimulating and sustaining root growth, maintaining soil biotic habitat, and minimizing soil deterioration, healthy soil ensures adequate retention and release of water and nutrients. Therefore, maintaining soil health is essential to agriculture to improve crop health, ensure food and nutrition, cope with biotic and abiotic environmental stressors, e.g., UV-B and high light, and enhance agricultural productivity and profitability. In addition to storing and cycling nutrients, high-quality soil also filters and buffers organic and inorganic substances, producing drinkable groundwater. Therefore, managing it must be a key component of strategy to handle plant health, coping with stress, population growth, and development pressures (Yadav et al., 2019a; Yadav et al., 2019b; Lingwan et al., 2024a; Job et al., 2025; Saini. et al.,2020; Koley et al.,2025).

This study provides a comparative overview of the growth, yield, and soil health of organic and conventional *Pant Basmati-1* systems, conducted in the northwestern plains (the rice-growing region of the Tarai) of India. Basmati rice is essential from an agronomic and economic point of view, as high-quality scented varieties like Pant Basmati-1 offer strong aroma and long grain length and fetch a high market price both within the country and abroad. Given the new generation consumers’ preference for quality, healthy sustainability, and environmental concern, and considering that the global organic food market is set to reach USD 380 billion by 2025 (Statista, 2024, Lingwan et al., 2016; Kushwaha et al., 2022), an analysis of organic viability for high-value crops becomes imperative. Various conventional management schemes have led to diminished soil fertility and are contrary to sustainable agricultural practices. Literature findings and data gaps in studies may contribute to the understanding of the growth parameters, yield components, and soil health of certain Basmati types such as *Pant Basmati-1*. In order to address the issue, this study determines the agronomic and biochemical characteristics in connection to different nutrient management strategies.

Compost and green manure (*Sesbania aculeata, Crotalaria juncea*) are examples of organic nutrient sources that do more than nourish the soil; they also maintain nutrient cycling and increase microbial diversity. That is based on current research, not merely theory (Texas A&M AgriLife Organic, 2024; FiBL, 2023). Recently, bio-augmentation-the addition of beneficial bacteria like *Pseudomonas fluorescens* and *Trichoderma harzianum*-has become popular. This increases microbial activity and accelerates decomposition (Singh et al., 2023). Conversely, traditional systems rely significantly on artificial inputs. Yes, you may immediately observe higher yields, but there is a cost: the variety of soil bacteria declines, heavy metals begin to accumulate, and the soil becomes worn out (Rahman et al., 2021; Sharma & Arora, 2023). This study examines crop growth, plant responses, yield, and soil health indicators. We are contrasting conventional synthetic fertilizers with organic methods based on green manuring. The objective is to provide policymakers, researchers, and farmers with reliable data to inform decisions on sustainable rice production.

Crucially, as demonstrated by long-term experiments such as the DOK experiment, our work confirms that organic systems need a transition period-typically more than two years-for soil biological equilibrium to establish (Mäder et al., 2002). However, organic plots showed encouraging long-term productivity indicators, including improved sustainability indices and potential economic viability. Furthermore, Pant Basmati-1’s aromatic characteristic, which results from the presence of 2-acetyl-1-pyrroline, remains a crucial marketing attribute (Kumar et al., 2023). For regions where consumer demand for residue-free rice is growing, our region-specific insights into organically managing this type are especially pertinent. With an emphasis on the true advantages and disadvantages of each rice variety, this study explores how organic rice farming functions in various geographical areas. It illustrates the difficult decisions between using conventional methods to increase harvests and switching to organic farming for environmental reasons. All of this contributes something tangible to the discussion about how to transition to more environmentally friendly agricultural practices and maximize inputs, particularly for important commodities like Basmati rice. Several studies have investigated how organic farming varies from conventional farming with respect to SOC content or stock. However, the results are difficult to generalize because they rely on variables like climate, the length of organic farming, and the precise management of both farming systems. However, most research has shown that organic farming increases the overall SOC concentration in top soils (Gattinger et al., 2012, Giannini et al., 2023). It is argued that the elevated SOC contents result from a high, or even disproportionately high, application of external OM inputs, which cannot be sustained with large-scale expansion of organic farming and only lead to carbon re-allocation rather than sequestration (Kirchmann et al., 2016, Gaudaré et al., 2023; Lingwan, & Yadav 2025).

The novelty of this work lies in its two-year, field-based comparative evaluation of organic and conventional nutrient management systems on a released Basmati rice variety (Pant Basmati 1) under subtropical Kharif conditions. Unlike many short-term or yield-only studies, this research integrates phenological stability, physiological responses, biomass partitioning, and soil quality indicators to assess system performance during the early phase of organic conversion. A key novel finding is the yield parity between organic and conventional systems despite consistently higher biomass production under inorganic fertilization, demonstrating that organic management can maintain productivity even during the initial years of transition. The study uniquely reports a significant increase in harvest index under organic management in the second year, indicating improved assimilate partitioning as soil fertility begins to stabilize, an aspect rarely quantified in Basmati rice under organic systems. Additionally, the work provides new insights into temporal shifts in growth dynamics (year- and stage-specific responses in plant height, LAI, chlorophyll content, and dry matter accumulation) rather than relying on single-stage observations organic management practices improves the soil health as compare to conventional management practices because of the Tarai soil are rich in organic matter content which improves the available macro and micronutrients and physical and biological properties. The finding that phonological traits remain unaffected by nutrient source further strengthens the understanding that nutrient management alters crop performance without disturbing crop duration in Basmati rice. Overall, this study contributes original evidence on the transition behaviour of organic nutrient management in aromatic rice, emphasizing that while organic systems may require more than two years to match conventional productivity fully, they offer environmentally sustainable yield stability with improved soil health and nutrient use efficiency, making them a viable alternative for sustainable Basmati rice cultivation.

## Material and Methods

### Experimental material and Weather Conditions

The experiment is located in A_4_ block of the Norman E. Borlaug Crop Research Centre of G. B. Pant University of Agriculture and Technology, Pantnagar, U. S. Nagar, Uttarakhand. (Fig. 1a) The experimental site lies in *the Tarai plains about 30 km southward of the foothill of the Shivalik range of the Himalayas at 290 N latitude, 790 29’ E longitude,* and an altitude of 243.8 m above the mean sea level. The study was conducted using *Pant Basmati-1*, a premium aromatic rice variety, with seeds obtained from the Department of Genetics and Plant Breeding, G.B. Pant University of Agriculture and Technology, Pantnagar. The Soil type is sandy loam with moderate fertility. A randomised block design (RBD) with three replications per treatment was employed.

**Figure 1.**
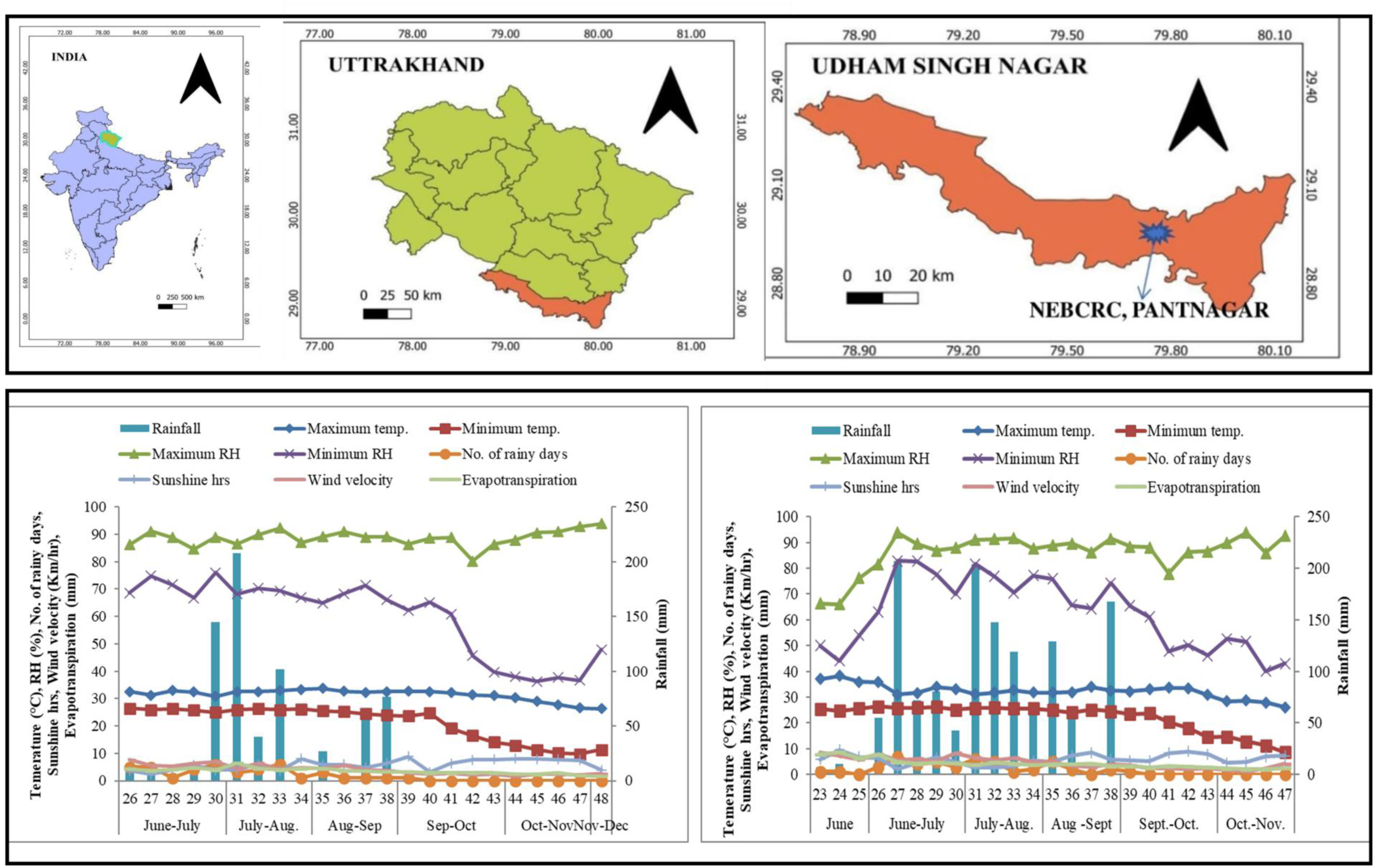
a. Map of the Location of experimental site 1 b) Standard meteorological week’s average weather data of season kharif, 2023,1 c) Standard meteorological week’s average weather data of season kharif, 2024

The experiment was carried out over two consecutive *Kharif* seasons (2023 and 2024) in two adjacent experimental plots, each measuring 450 m². One plot followed conventional (inorganic) management, receiving a fertilizer dose of 120:60:40 kg N:P₂O₅:K₂O per hectare, while the other was managed under organic practices, with Sesbania rostrata sown as green manure and incorporated at 55 days after sowing (DAS), 1 day before transplanting. Seedlings were raised in a dry bed nursery and transplanted at 21 days old with a spacing of 20 cm × 10 cm (row × plant). All standard agronomic practices recommended for the region were followed. The crop was harvested manually at 90% panicle maturity, followed by threshing and sun drying. Grain yield was determined as weight per plot and converted into tonnes per hectare (t/ha).

Seasonal weather conditions during 2023 and 2024 were recorded, including total rainfall, average daily temperature, relative humidity, number of rainy days, wind velocity, sunshine, and evapotranspiration (Fig.1b, 1c).

### Observations recorded

#### Parameters of plant growth

##### Plant height, Number of tillers and Dry matter accumulation

Measured from the soil surface to the tip of the tallest leaf on three randomly selected plants at 30, 60, and 90 days after transplanting (DAT) using a meter scale. The number of tillers was counted per square meter in each replication at 30, 60, and 90 DAT. Dry matter accumulation was analysed from Plants sampled at 30, 60, and 90 DAT, which were oven-dried at 65 ± 5°C for 48 hours, and the weight was recorded in g/m².

##### Leaf area index

LAI determined at the flowering stage from three hills per replication using the formula: LAI = (A× Number of leaves per hill) / 200

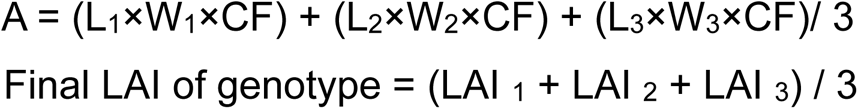

Where,L_1_ = Average length of 3 large leaves of hill, W_1_ = Average width of 3 large leaves of hill, L_2_ = Average length of 3 medium leaves of hill, W_2_ =Average width of 3 medium leaves of hill, L_3_ = Average length of 3 small leaves of hill, W_3_= Average width of 3 small leaves of hill, CF = Correction factor (0.75)

#### Phonological studies

##### Days to panicle Initiation, Days to flowering and Days to maturity

Beginning at 30 DAT, tillers were sampled every three days. Panicle initiation was identified through microscopic dissection using a sewing needle and noted upon the appearance of white hairs on the young panicle. Days to flowering were recorded when 50% of the panicles had fully emerged, and spikelets began anthesis, with visible stigma and anther extrusion. Days to maturity were defined when 90% of the grains on selected panicles turned straw-coloured, and the biting test confirmed grain hardness.

##### Chlorophyll content of leaves

Chlorophyll a, b, and total chlorophyll contents were estimated following Arnon’s method (1949) (Pant et al., 2023). At flowering, 50 mg of fresh leaf tissue was incubated in 7 mL DMSO at 65°C for 3 hours. After cooling, an additional 3 mL DMSO was added, and absorbance was recorded at 663 nm and 645 nm using a spectrophotometer. Concentrations were calculated using the following equations:

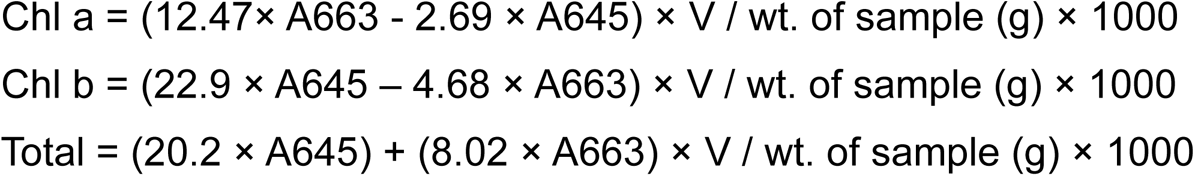

Where, A663 = Absorbance at 663 nm, A645 = Absorbance at 645 nm, V = Volume of DMSO used, W = Weight of leaves in gram

#### Parameters of yield at harvest

##### Total biological yield, Grain yield (t/ha) and Harvest index (%)

For the total biological yield biomass from a 1 m² area in each replication was uprooted, sun-dried, and weighed. Grain yield (t/ha) was calculated fromGrain output from each plot, which was weighed and converted to tonnes per hectare. Harvest index (%) is calculated as the yield of the plant parts of economic interest (economic yield) as a percentage of total biological yield in terms of dry matter.

##### Collection of soil samples and processing

A composite soil sample from each plot was collected after harvesting of rice crop in the year 2023-2024. In each plot, soil was collected randomly from five points and combined into a single sample. The soil sample from each plot was stored at a low temperature (0-40 C) in the refrigerator for the study of biological properties. The remaining portion of the soil sample was air-dried in the shade, passed through a 2 mm sieve, and stored for estimating its chemical and physical properties.

#### Available nitrogen observations

Alkaline potassium permanganate method was used for the determination of available nitrogen (Subbiah and Asija, 1956). Available phosphorus in soil was estimated by using sodium bicarbonate extractant (0.5 M NaHCO_3_) adjusted to pH 8.5 (Olsen *et al*., 1954). Available K was determined by neutral ammonium acetate (1N NH_4_OAc; pH 7) method described by Black, (1965). The modified Walkley and Black (1934) method was used to determine organic carbon, as described by Jackson (1967). Available Zn, Cu, Mn, and Fe were estimated by the method developed by Lindsay and Norvell (1978) using an atomic absorption spectrophotometer. Water holding capacity was determined with the help of Hilgard apparatus (Piper, 1950). Modified Yoder’s wet sieving method (Van Bavel, 1953) was used to determine Aggregate stability. The dehydrogenase activity of soil was determined using the method listed by Casida *et al*. (1964). The total phosphatase activity (acid and alkaline phosphatases) was determined according to the method of Tabatabai and Bremner (1969).

#### Statistical analysis

All recorded data were analysed using IBM SPSS Statistics. Initially, a paired t-test was applied to compare organic and inorganic treatments. Normality (Shapiro-Wilk test) and homogeneity of variance (Levene’s test) were verified before applying parametric tests. To assess treatment effects over time, a two-way factorial ANOVA was also employed, with year (2023, 2024) and treatment (organic, inorganic) as fixed factors (Shree et al., 2019). The significance of main effects and interaction terms (year × treatment) was tested to account for inter-annual variability and enhance statistical robustness. Significance was determined at p ≤ 0.05.

## Results

### Phonological Parameters

Phonological traits such as days to panicle initiation, flowering, and maturity in *Pant Basmati 1* were marginally influenced by nutrient management. Across both Kharif seasons (2023 and 2024), differences in days to panicle initiation and flowering between organic and conventional systems were statistically insignificant. On average, panicle initiation occurred between 58-62 days, and flowering between 84-88 days after transplanting. Similarly, maturity ranged from 106 to 111 days, with no consistent trend favouring either system (Fig. 2a). These results suggest that nutrient source did not significantly alter the phenological timing of the crop.

**Figure 2.**
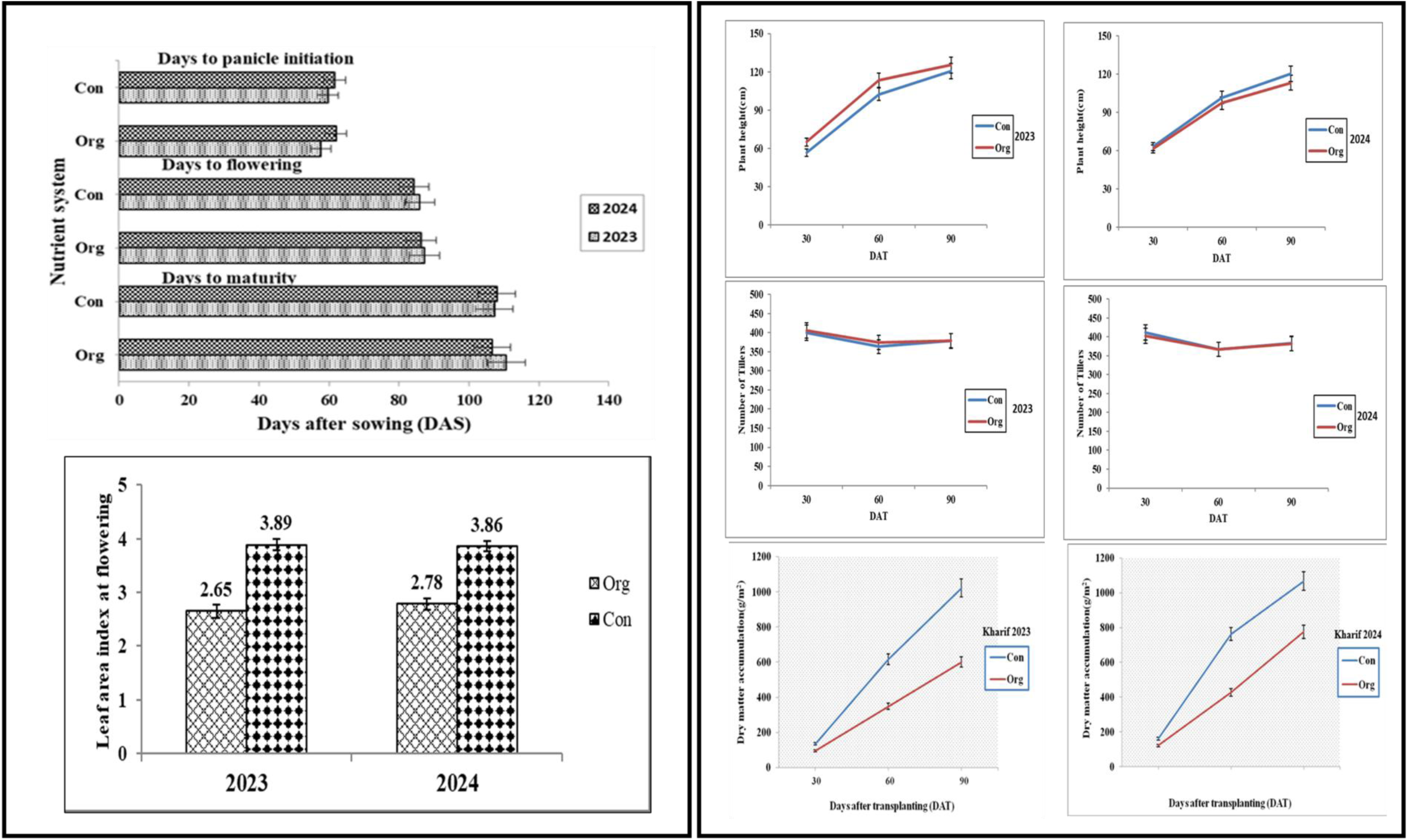
a) Effect of inorganic and organic sources of nutrients on days to Panicle initiation, days to flowering, and days to maturity in rice variety Pant Basmati 1 during Kharif 2023 and 2024. (2b). Effect of inorganic and organic sources of nutrients on plant height of rice variety Pant Basmati 1 at 30, 60, and 90 DAT during Kharif, 2023 & 2024. (2c). Effect of inorganic and organic sources of nutrients on the number of tillers of rice variety Pant Basmati 1 at 30, 60, and 90 DAT during Kharif, 2023 & 2024. 2d). Effect of inorganic and organic sources of nutrients on leaf area index at flowering in rice variety Pant Basmati 1 (Kharif 2023 and 2024). 2e) Effect of inorganic and organic sources of nutrients on dry matter accumulation of rice variety Pant Basmati 1 at 30, 60, and 90 DAT, Kharif 2023& 2024.

### Morphological Parameters

The influence of different nutrient sources on key morphological parameters, including plant height, number of tillers per plant, and leaf area index (LAI), was systematically assessed to evaluate the impact of organic and inorganic nutrient applications on rice plant growth. These metrics are important markers of overall crop development, biomass accumulation, and plant vigor. The study aims to shed light on how well organic and inorganic fertilization optimize rice growth and productivity by examining differences in these characteristics across various nutrient regimes.Plant height differed across seasons and growth stages (Fig. 2b). In 2023, organic plots exhibited significantly taller plants at 30 DAT, although differences evened out by 60 and 90 DAT. Conversely, in 2024, inorganic treatments led to significantly taller plants at 90 DAT, suggesting sustained nutrient release from mineral fertilizers at later stages.

Tiller counts remained statistically similar across both systems and years (Fig. 2c). Tillers ranged from 363 to 411 m², indicating that both nutrient sources supported adequate vegetative proliferation, with organic inputs performing comparably to conventional fertilizers. LAI was consistently higher in the inorganic system across both years (Fig. 2d), reaching 3.86-3.89, compared to 2.65-2.78 under organic management. This highlights the immediate nutrient availability under inorganic regimes, promoting greater canopy expansion. Dry matter accumulation (Fig. 2 and 4) was significantly greater in the inorganic system at all observed stages. The trend was consistent across years, underlining the role of mineral fertilizers in promoting continuous biomass development.

### Physiological and Biochemical Parameters

#### Chlorophyll ‘a’, ‘b’, and Total Chlorophyll

While not statistically significant, chlorophyll ‘a’ and ‘b’ contents peaked at 60 DAT in the organic system (Fig.3a, 3b, 3c), suggesting a possible benefit of organic inputs in maintaining photosynthetic pigments during critical growth phases. Biological yield was higher in the inorganic system (11.60 t/ha in 2023; 10.63 t/ha in 2024), although differences were not statistically significant (Fig. 11). Grain yield also showed no significant variation across treatments (Fig. 4a), pointing to the potential of organic systems to maintain yield parity with conventional methods over time. Harvest index remained unaffected in the first year. However, it increased significantly under organic management in the second year (Fig. 4b). This may indicate improved partitioning efficiency in organic systems as soil fertility stabilizes.

#### Available Macronutrient (Nitrogen, Phosphorous and Potassium)

The available macronutrients, i.e. nitrogen, phosphorus, and potassium, were observed highest under the organically treated plot as compared to the conventional plot **(Table 1).** In 2023, organic management supplied a greater amount of available nitrogen (159.20 kg ha⁻¹) than conventional management (152.46 kg ha⁻¹). Similar outcomes were observed in 2024. In 2023, organic plots had higher available P (16.84 kg ha⁻¹) compared to conventional plots (14.94 kg ha⁻¹). In 2024, the trend continued, with 17.51 kg ha⁻¹ in organic plots and 15.60 kg ha⁻¹ in conventional plots. Organically treated plots had higher available K levels (153.20 and 155.20 kg ha⁻¹, respectively) than conventional plots (148.80 and 149.46 kg ha⁻¹, respectively) for both years.

**Table 1.** Effect of organic and conventional management practices on soil macronutrients and organic carbon during *kharif* season of 2023 and 2024

| Soil Properties | Year | Treatment Means (±SD) | t-value | p-value | Significance |
| --- | --- | --- | --- | --- | --- |
| Available N ($\text{kg ha}^{-1}$) | 2023 | Org $159.20 \pm 3.48$<br>Con $152.46 \pm 2.59$ | 4.06 | 0.014 | * |
| | 2024 | Org $161.20 \pm 2.59$<br>Con $154.46 \pm 4.06$ | 2.75 | 0.05 | * |
| Available P ($\text{kg ha}^{-1}$) | 2023 | Org $16.84 \pm 1.55$<br>Con $14.94 \pm 0.66$ | 2.82 | 0.047 | * |
| | 2024 | Org $17.51 \pm 1.51$<br>Con $15.60 \pm 0.54$ | 2.66 | 0.056 | NS |
| Available K ($\text{kg ha}^{-1}$) | 2023 | Org $153.20 \pm 5.16$<br>Con $148.80 \pm 7.70$ | 1.55 | 0.19 | NS |
|  | 2024 | Org155.20 ± 4.96<br>Con149.46 ± 10.47 | 1.63 | 0.18 | NS |
| Organic C (%) | 2023 | Org 0.87 ± 0.03<br>Con 0.81 ± 0.03 | 3.18 | 0.033 | * |
|  | 2024 | Org 0.93 ± 0.05<br>Con 0.82 ± 0.03 | 3.26 | 0.03 | * |

#### Organic Carbon (%) and available Micronutrients (Zinc, Iron, Manganese and Copper)

In both years, the Organic carbon content was significantly higher under organically (0.87 and 0.93 % respectively) treated plots compared to the inorganically (0.81 and 0.82 % respectively) **(Table 1)**

The available micronutrients, i.e., Zinc, Iron, Manganese, and Copper, were received higher with organic practices as compared to inorganic (Table 2). Zinc was slightly higher in organic plots in both years. In 2023, the organically treated plot had higher Zn (1.14 mg kg⁻¹) than the conventional method (0.97 mg kg⁻¹). Similar trends were also observed in 2024. Fe showed consistently higher availability under organic management (19.48 mg kg⁻¹ in 2023 and 20.28 mg kg⁻¹ in 2024) than under conventional management (17.21 and 18.21 mg kg⁻¹). Available Mn was tuned higher under the organically treated plot (5.42 mg kg⁻¹ in 2023 and 5.86 mg kg⁻¹ in 2024) as compared to the conventional (4.52 and 5.18 mg kg⁻¹). Copper was also attained at higher levels under organic practices (3.53 mg kg⁻¹ and 3.98 mg kg⁻¹ respectively), as compared to conventional practices (2.52 and 3.25 mg kg⁻¹ respectively).

**Table 2.** Effect of organic and conventional management practices on soil micronutrients during *kharif* season of 2023 and 2024

| Soil Properties | Year | Treatment Means (±SD) | t-value | p-value | Significance |
| --- | --- | --- | --- | --- | --- |
| Zn (mg kg<sup>-1</sup>) | 2023 | Org 1.14 ± 0.05<br>Con 0.97 ± 0.15 | 3.08 | 0.036 | * |
|  | 2024 | Org 1.18 ± 0.06<br>Con 1.03 ± 0.10 | 2.5 | 0.067 | NS |
| Fe (mg kg<sup>-1</sup>) | 2023 | Org 19.48 ± 2.08<br>Con 17.21 ± 0.94 | 4.21 | 0.013 | * |
|  | 2024 | Org 20.28 ± 1.04<br>Con 18.21 ± 0.07 | 2.83 | 0.047 | * |
| Mn (mg kg<sup>-1</sup>) | 2023 | Org 5.42 ± 1.01<br>Con 4.52 ± 0.51 | 3.45 | 0.025 | * |
|  | 2024 | Org 5.86 ± 0.85<br>Con 5.18 ± 0.72 | 2.61 | 0.059 | NS |
| Cu (mg kg<sup>-1</sup>) | 2023 | Org 3.53 ± 0.49<br>Con 2.52 ± 0.85 | 3.09 | 0.035 | * |
|  | 2024 | Org 3.98 ± 0.58<br>Con 3.25 ± 0.42 | 2.71 | 0.053 | NS |

#### Water-Holding Capacity (%) and Aggregate Stability (%)

Organic treatment significantly enhanced WHC, with results of 59.80% in 2023 and 61.13% in 2024, compared to 53.15% and 54.49% with conventional treatment (Table 3). Improved WHC is associated with increased organic matter, aggregation, and stability of pore space. Aggregate stability improved significantly in organic plots (19.72% in 2023 and 20.44% in 2024) compared to conventional plots (17.09% and 18.87%).

**Table 3.** Effect of organic and conventional management practices on soil physical and biological properties during the *kharif* season of 2023 and 2024

| Soil Properties | Year | Treatment Means<br>(±SD) | t-value | p-value | Significance |
| --- | --- | --- | --- | --- | --- |
| WHC (%) | 2023 | Org 59.80 ± 0.55<br>Con 53.15 ± 2.60 | 9.05 | 0.001 | ** |
| | 2024 | Org 61.13 ± 1.01<br>Con 54.49 ± 2.06 | 6.13 | 0.003 | ** |
| Aggregate stability (%) | 2023 | Org 19.72 ± 0.61<br>Con 17.09 ± 1.09 | 7.12 | 0.002 | ** |
| | 2024 | Org 20.44 ± 0.48<br>Con 18.87 ± 2.07 | 3.41 | 0.025 | * |
| Alkaline phosphatase<br>($\mu\text{g PNP g}^{-1} \text{ h}^{-1}$) | 2023 | Org 42.46 ± 2.52<br>Con 38.15 ± 2.53 | 2.71 | 0.053 | NS |
| | 2024 | Org 43.45 ± 1.72<br>Con 39.15 ± 2.87 | 2.95 | 0.042 | * |
| Acid phosphatase ($\mu\text{g}$<br>$\text{PNP g}^{-1} \text{ h}^{-1}$) | 2023 | Org 48.13 ± 2.55<br>Con 40.79 ± 1.49 | 5.44 | 0.006 | ** |
| | 2024 | Org 50.80 ± 1.05<br>Con 42.45 ± 2.19 | 4.02 | 0.015 | * |
| Dehydrogenase (µg<br>TPF g<sup>-1</sup> day<sup>-1</sup>) | 2023 | Org 231.82 ± 2.94<br>Con 226.19 ± 1.69 | 2.66 | 0.057 | NS |
|  | 2024 | Org 235.48 ± 3.54<br>Con 228.18 ± 1.64 | 3.07 | 0.036 | * |

#### Alkaline, acid Phosphatase and Dehydrogenase Activity (DHA)

During the year 2023, the organically treated plots had higher alkaline and acid phosphatase activity (42.46 and 48.13 µg PNP g⁻¹ soil h⁻¹) than conventional plots (38.15 and 39.15 PNP g⁻¹ soil h⁻¹). However, the difference was not statistically significant **(Table 3).** A similar pattern was observed in 2024, but it had a significant effect. During both years, the DHA levels were higher under organic management techniques compared to conventional methods **(Table 3)**. In 2023, organic and conventional practices exhibited no significant effect on DHA (231.82 vs 226.19 µg TPF g⁻¹ day⁻¹), whereas, in 2024 (235.48 vs 228.19 µg TPF g⁻¹ day⁻¹), the organic and conventional methods showed a significant impact. This pattern suggests rising microbial oxidative activity and respiration under organic management.

## Discussion

The differential response of *Pant Basmati 1* to organic and inorganic nutrient sources highlights key physiological and agronomic dynamics. Plant height and leaf area index (LAI)-both vital to crop productivity-were consistently higher under inorganic management, especially during later growth stages. Gebrekidan and Seyoum (2006) also noted that mineral fertilizers significantly shape canopy growth and help plants accumulate more biomass. The increased LAI is evident in the inorganic system (Fig. 2), improving photosynthetic efficiency by allowing the crop to receive more sunlight. It is not surprising that the inorganic plots had a higher biological yield (Fig. 3, 4) because LAI and dry matter both closely correlate with grain production (Oo et al., 2010; Yadav et al., 2020). The harvest index (Fig.4) increased with the organic treatment during the second year, which is an intriguing turn of events. This suggests that the plant will be more efficient in allocating its resources and utilizing nutrients, particularly once the soil settles into its new organic groove-something that Mäder et al. (2002) also observed. Even more intriguingly, the organic plots’ levels of chlorophyll “a” and “b” peaked at 60 DAT (Fig. 2, 3, 4). Therefore, organic crops appear to increase photosynthetic efficiency, even though they may begin more slowly. This is likely related to a continual flow of nutrients and more active soil bacteria, as detailed by Siavoshi and Laware (2013). The fact that there was no variation in grain yields between treatments (see Fig.4) goes to demonstrate that organic systems can compete with conventional ones following the changeover and can be adopted in the future to other abiotic stresses like UV-B or high light (Job et al., 2022; Lingwan et al., 2024b; Thakur et al., 2025; Lingwan et al., 2024c). This is consistent with ongoing research, such as the DOK experiment. It is not easy to get there, however. During the transition, farmers must contend with increased pest issues, unpredictable weather, and delays in nutrient supplies. These time-related hurdles make it clear why we need long-term field trials and studies that really track how benefits build up over time.

**Fig. 3.**
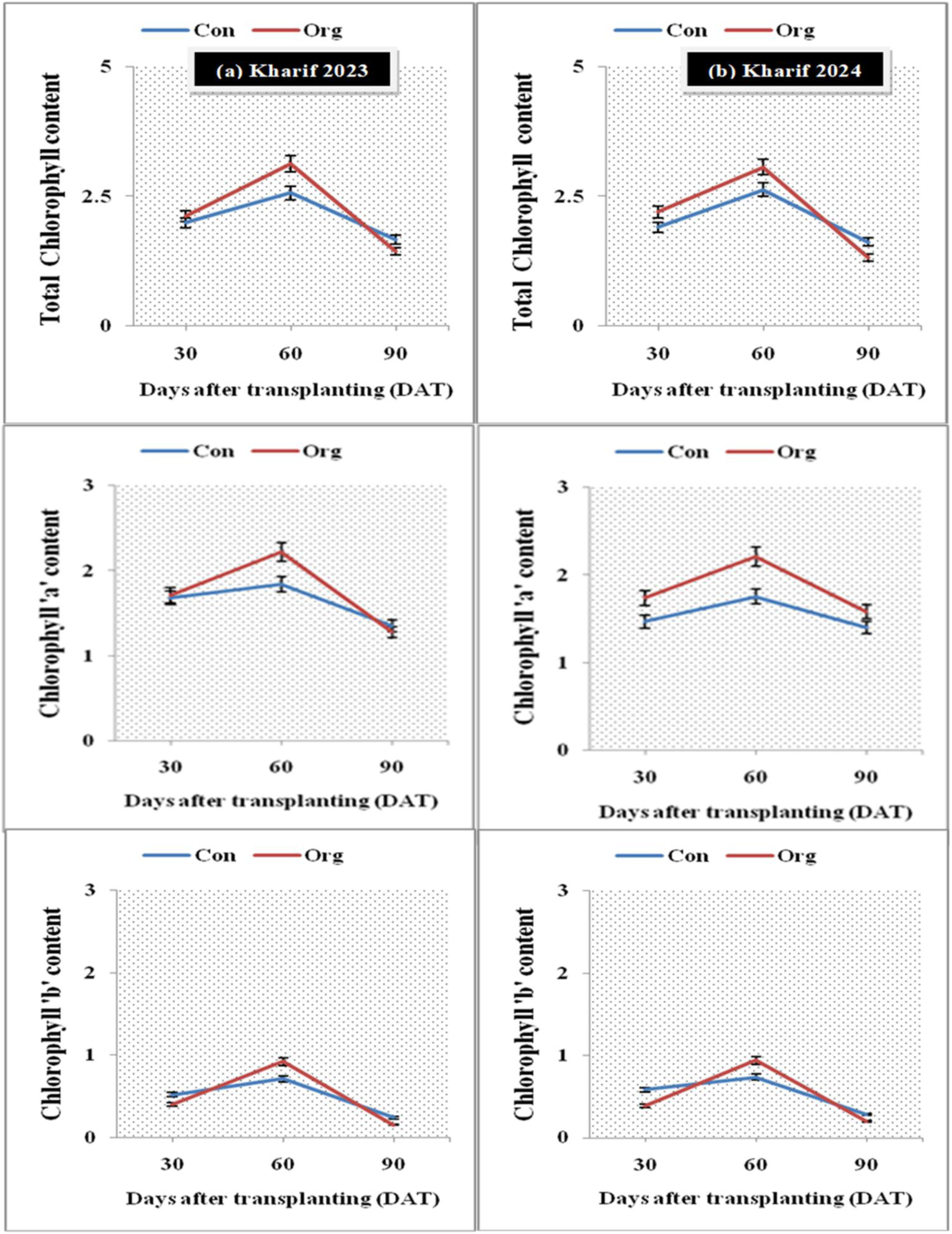
a) Effect of inorganic and organic sources of nutrients on total chlorophyll content (mg/g fresh wt.) of rice variety Pant Basmati 1 at 30, 60 and 90 DAT, Kharif 2023 (a) and 2024 (b), 3b) Effect of inorganic and organic sources of nutrients on chlorophyll ‘a’content (mg/g fresh wt.) of rice variety Pant Basmati 1 at 30, 60 and 90 DAT, Kharif 2023 (a) and 2024 (b) 3c). Effect of inorganic and organic sources of nutrients on chlorophyll ‘b’ content (mg/g fresh wt.) of rice variety Pant Basmati 1 at 30, 60, and 90 DAT, Kharif 2023 (a) and 2024(b)

**Fig. 4.**
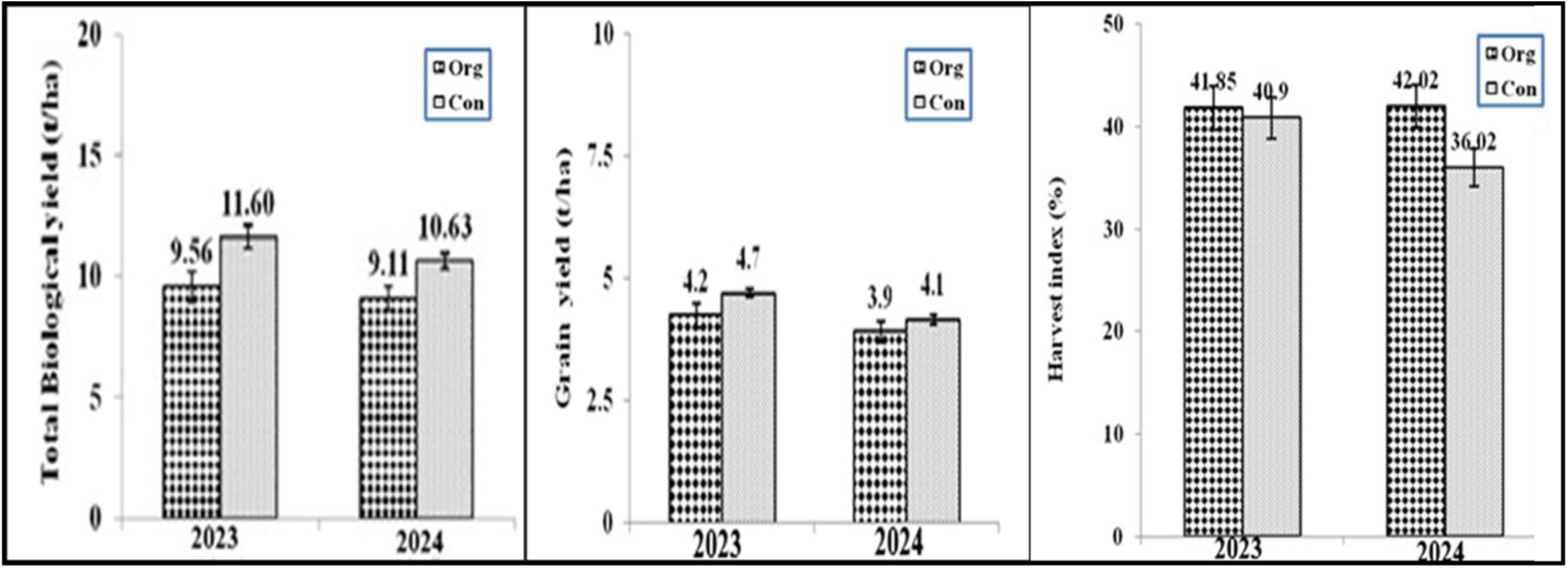
a. Effect of inorganic and organic sources of nutrients on total biological yield (t/ha) of rice variety Pant Basmati 1, Kharif 2023 and 2024. 4b). Effect of inorganic and organic sources of nutrients on grain yield (t/ha) of rice variety Pant Basmati 1, Kharif 2023 and 2024. 4c). Effect of inorganic and organic sources of nutrients on harvest index (%) in rice variety Pant Basmati 1 (Kharif 2023 and 2024)

This is supported by research from developing regions. According to Zúniga-González and colleagues (2022), field schools in Nicaragua that taught sustainable practices increased technical efficiency. The bioeconomic benefits of environmental sustainability in organic systems were previously highlighted by Blanco and Zúniga-González (2013). Our findings support the same conclusion: although organic farms may initially produce lower yields, the long-term financial and environmental benefits make organic a viable choice. Looking ahead, there is room to dig deeper. Future research should examine system productivity, multi-omics, metabolic modeling integrating data envelopment analysis (Zúniga-González et al., 2024; Lingwan et al., 2025a; Ali et al., 2025b; Lingwan et al., 2025b; Koley et al., 2024; Lingwan et al., 2025c), and factor in both bioeconomic and climate resilience metrics (Zúniga-González, 2023; Mohanasundaram et al., 2021). The study’s findings clearly demonstrate that organic farming methods greatly enhanced soil health indicators, confirming the initial perspective. Compared to conventional farms, organic farms exhibited more soil organic matter, higher nutrient content, higher WHC and aggregate stability, and higher enzymatic activity. These improvements can be associated with organic farming’s use of natural inputs, which increase soil structure and nutrient cycling. However, because conventional agricultural methods rely on synthetic fertilizers, soil minerals are depleted more quickly. Even though these chemical inputs release nutrients quickly, they frequently cause long-term soil deterioration, decreased microbial activity, and acidification. Over time, even greater inputs of synthetic fertilizers may be required due to the faster nutrient loss in conventional systems, which would continue the cycle of declining soil health.

The mineralization of nitrogen and increased growth of soil microorganisms, which turn organically bound nitrogen into inorganic form, may be liable for the increase in available nitrogen content in soil caused by the application of organic matter. Enhanced nitrogen availability in organic farms, as noted by (Meena et al., 2020; Bhanuvally et al., 2024; Panghate et al., 2020). As organic matter is used, an accumulation of available P may be ascribed to the release of organic acids during decomposition, which in turn facilitates the release of native phosphorus by the solubilizing activity of these acids. Additionally, organic matter binds sesquioxides, making them inactive and lowering the soil’s ability to fix phosphate, which ultimately assists in the release of an adequate quantity of plant-available P. Addition of organic matter improves the available P, as also reported by (Subehia & Sepehya, 2012; Dhaliwal et al., 2015; Cullen et al., 2025).In addition to the direct supply of K to the soil, organic matter may be attributed to the decrease in K fixation and release of K resulting from the interaction of organic matter with clays ( Subehia & Sepehya, 2012; Dhaliwal et al., 2015). The use of organic matter may have promoted soil organic carbon through increased root biomass and functioned as a binding agent, promoting aggregate formation. The microbial degradation of the organic residues added to the soil emits various organic products of decay, such as polysaccharides, which act as beneficial cementing agents in the formation of large, stable aggregates that enhance soil aggregate stability. Similar work was also reported by Sihi et al. (2017) and Williams et al. (2017). The higher level of soil organic carbon with the addition of organic matter may be due to enhanced root growth, resulting in more organic residue in soil, which, after decomposition, might have increased the soil organic carbon content. These results conform to the findings of Singh *et al*. (1999) in Mollisols at Pantnagar.

The release of chelating agents from the decomposition of organic matter and the accessibility of micronutrients through organic matter may have prevented their precipitation, oxidation, and leaching. The addition of micronutrients through organic matter and the release of chelating agents from organic matter decomposition have been attributed to the increase in the available Zn, Fe, Mn, and Cu status of soils in organically treated plots. These factors may have prevented micronutrients from precipitating, oxidizing, and leaching. The lack of micronutrient replenishment with chemical fertilizers is the reason for the decrease in micronutrient levels in inorganic treatments (Lingwan et al., 2022; Lingwan et al., 2023). As organic matter is added, the soil becomes more porous, and stable aggregates form due to increased microbial activity that produces polysaccharides, glomalin, etc., thereby increasing the soil’s water-holding capacity. (Bhatt et al., 2019; Kumar and Singh, 2010; Kharche et al., 2013; Jaroensuk et al., 2025). These stable aggregates have greater potential to retain water, thereby increasing the soil’s water-holding capacity (Suja et al., 2017; Sarkar et al., 2024). Due to the increased availability of substrate from organic manure, which leads to high microbial activity and the release of these enzymes into the soil, the population of microorganisms may be responsible for the increase in acid and alkaline phosphatase and Dehydrogenase activity in the soil. An increase in the microbial population may also be responsible for the rise in acid, alkaline, and DHA activity in organically treated plots (Jha et al., 1992; Lugato et el., 2018; Anand et al., 2015; Rathore et al., 2024)

## Conclusion

According to the study, organic basmati rice farming-especially when using green manure like Sesbania rostrata-pays off in the long run, whereas conventional systems provide a short-term productivity boost. The harvest index increases, the soil becomes healthier, and nutrients are delivered more steadily. Yields are roughly the same in the second year, but it takes more than 2 years for the actual advantages of becoming organic to become apparent. Consistent increases in the harvest index and chlorophyll support this trend. All of this suggests that maintaining organic practices throughout time is crucial for basmati rice. Although there are initial challenges, organic farming increases soil fertility, reduces environmental hazards, and maintains sustainable food production. Ultimately, it is a sustainable option that aligns with global sustainability objectives and serves as a good alternative to traditional farming.

## Acknowledgment

I want to thank G.B. Pant University of Agriculture and Technology and D.S.B. Campus, Kumaun University, Nainital. They provided the facilities and support I needed to complete this research on organic and conventional crop production. I am also grateful to the faculty, technical staff, and fellow researchers who offered their guidance and help along the way.

## Authors Contributions

HCJ: Writing – original draft, Writing – review & editing; BP, MKB, MS: Supervision, Writing – review & editing*, SK: Writing – review & editing, Supervision.

